# The *Drosophila melanogaster* enzyme glycerol-3-phosphate dehydrogenase 1 is required for oogenesis, embryonic development, and amino acid homeostasis

**DOI:** 10.1101/2022.02.15.480380

**Authors:** Madhulika Rai, Sarah M. Carter, Shefali A. Shefali, Nader H. Mahmoudzadeh, Robert Pepin, Jason M. Tennessen

## Abstract

As the fruit fly, *Drosophila melanogaster*, progresses from one life stage to the next, many of the enzymes that compose intermediary metabolism undergo substantial changes in both expression and activity. These predictable shifts in metabolic flux allow the fly meet stage-specific requirements for energy production and biosynthesis. In this regard, the enzyme Glyerol-3-phosphate dehydrogenase (GPDH1) has been the focus of biochemical genetics studies for several decades, and as a result, is one of the most well characterized *Drosophila* enzymes. Among the findings of these earlier studies is that GPDH1 acts throughout the fly lifecycle to promote mitochondrial energy production and triglyceride accumulation while also serving a key role in maintaining redox balance. Here we expand upon the known roles of GPDH1 during fly development by examining how depletion of both the maternal and zygotic pools of this enzyme influences development, metabolism, and viability. Our findings not only confirm previous observations that *Gpdh1* mutants exhibit defects in larval development, lifespan, and fat storage but also reveal that GPDH1 serves essential roles in oogenesis and embryogenesis. Moreover, metabolomics analysis reveals that a *Gpdh1* mutant stock maintained in a homozygous state exhibits larval metabolic defects that significantly differ from those observed in the F1 mutant generation. Overall, our findings highlight unappreciated roles for GPDH1 in early development and uncover previously undescribed metabolic adaptations that could allow flies to survive loss of this key enzyme.

## INTRODUCTION

The *Drosophila* enzyme GPDH1 (encoded by FBgn0001128) is an ideal model for understanding how metabolism adapts to the biosynthetic and energetic requirements of animal development. Although the reaction catalyzed by GPDH1 is relatively simple (Figure 1), the activity and purpose of the enzyme varies significantly during development. For example, GPDH1 is highly expressed in both the larval fat body and adult flight muscle but has opposite functions within the two tissues. The larval fat body displays high levels of GPDH1 activity and relies on this enzyme to generate glycerol-3-phosphate (G3P; Figure 1), which is used in triglyceride synthesis (TAG) (Rechsteiner 1970; Sullivan *et al*. 1983; Merritt *et al*. 2006; Li *et al*. 2019). Meanwhile, GPDH1 in adult flight muscle functions in conjunction with the mitochondrial enzyme GPO1 to shuttle electrons into the electron transport chain for ATP synthesis (Sacktor and Dick 1962; O’brien and macIntyre 1972b; O’brien and macIntyre 1972a; Wojtas *et al*. 1997; Merritt *et al*. 2006), thus supporting the intense energy demands of insect flight (Figure 1).

**Figure 1.**
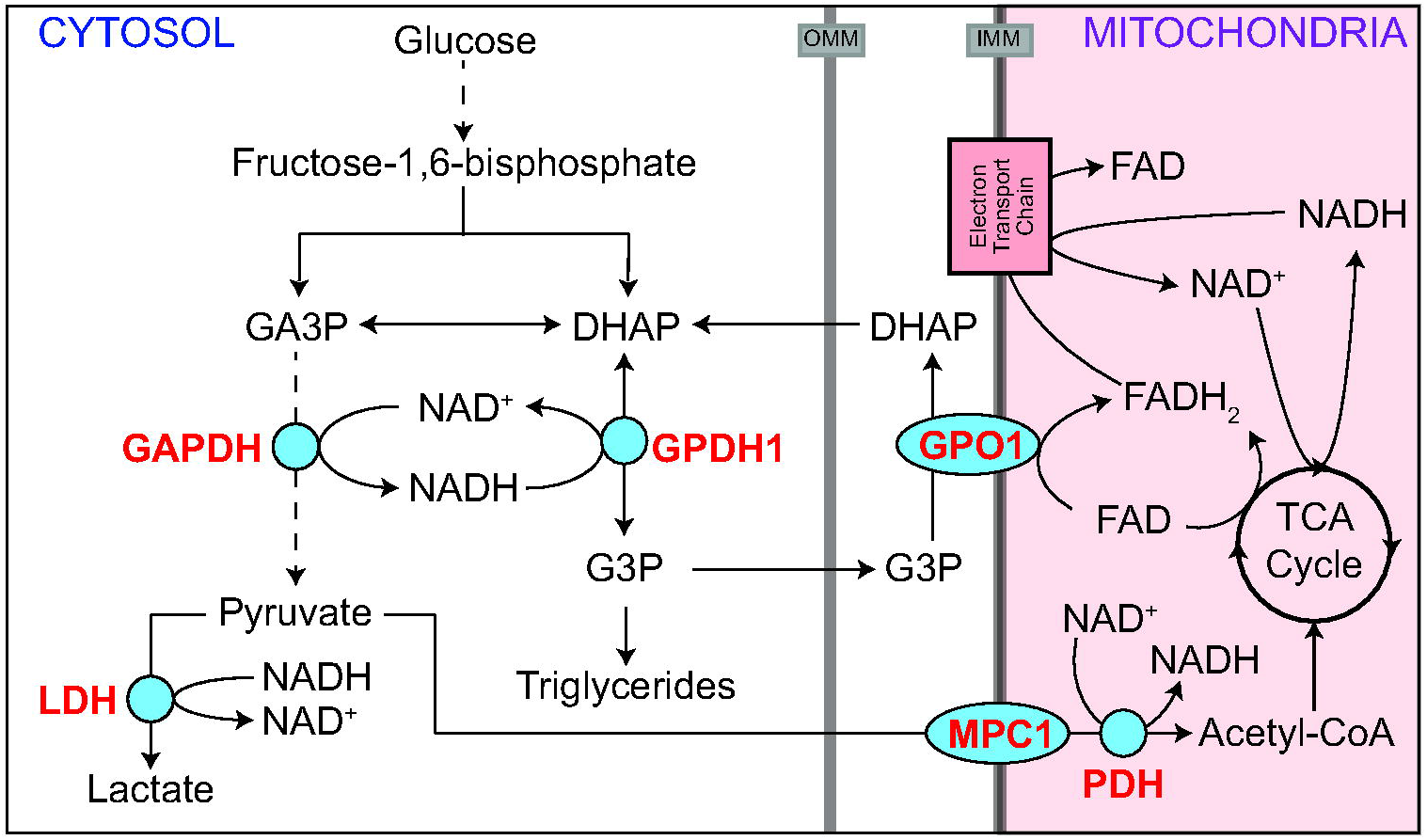
GPDH1 promotes cytosolic redox balance, ATP production, and TAG accumulation. A schematic diagram illustrating the role of GPDH1 in central carbon metabolism. GPDH1 relies on the cofactor NAD^+^/NADH to interconvert the glycolytic intermediate dihydroxyacetone phosphate (DHAP) to glycerol 3-phosphate (G3P). In *Drosophila* larvae, the GPDH1-dependent conversion of DHAP to G3P functions in parallel with Lactate Dehydrogenase (LDH) to maintain redox balance. G3P is used as a precursor for TAG synthesis and functions in the G3P electron shuttle to transfer reducing equivalents to the electron transport chain. In adult flight muscle, GPDH1 function in conjunction with GPO1 and the tricarboxylic acid (TCA) cycle to generate the ATP required for flight. Abbreviations: dihydroxyacetone phosphate (DHAP), glyceraldehyde-3-phosphate (GA3P), glycerol-3-phosphate (G3P), glycerol-3-phosphate dehydrogenase (GPDH1), glycerophosphate oxidase 1 (GPO1), lactate dehydrogenase (LDH), mitochondrial pyruvate carrier 1 (MPC1), pyruvate dehydrogenase (PDH), tricarboxylic acid (TCA).

The distinct functions of GPDH1 within the larval fat body and adult flight muscles illustrate why this enzyme serves as a model for understanding how metabolism is coordinately regulated in the context of animal growth, development, and physiology. Nearly fifty years of intensive biochemical and genetic studies revealed that GPDH1 kinetics, stability, and physical characteristics vary dramatically between the larval and adult enzyme pool and that GPDH1 activity fluctuates as a function of developmental time, with activity levels peaking in L3 larvae and in adults (Wright and Shaw 1969; Rechsteiner 1970; O’brien and macIntyre 1972a; Sullivan *et al*. 1983). These biochemical studies, coupled with decades of genetic analysis (O’brien and macIntyre 1972b; O’brien and Shimada 1974; Bewley *et al*. 1980; Kotarski *et al*. 1983; Burkhart *et al*. 1984; Davis and macIntyre 1988; Gibson *et al*. 1991; Yamaguchi *et al*. 1994; Merritt *et al*. 2006; Carmon *et al*. 2010; Li *et al*. 2019), make GPDH1 among the most intensively studied enzymes in developmental biology and provide insight towards how normal animal development relies on dramatic changes in enzyme activity.

Despite *Gpdh1* serving as the subject of dozens of biochemical genetic studies, one pool of GPDH1 remains largely overlooked. *Drosophila* embryos contain a significant amount of maternally-loaded GPDH1 that potentially persists into larval development (Wright and Shaw 1969; Wright and Shaw 1970; Casas-Vila *et al*. 2017). Yet, nearly all *Gpdh1* mutant studies only examine zygotic mutants, raising the possibility that novel GPDH1 functions could be discovered by examining mutants lacking the maternal enzyme pool. Here we address this possibility by examining the metabolic and developmental defects displayed by maternal-zygotic (M/Z) *Gpdh1* mutants. Our analysis reveals GPDH1 is not only required for normal oocyte development but also that loss of the maternal GPDH1 pool leads to a significant embryonic lethal phenotype. In contrast, among those *Gpdh1 M/Z* mutants that survive the embryonic lethal phase, post-embryonic development and adult longevity appears similar between zygotic and M/Z *Gpdh1* mutants. During our studies, however, we made the unexpected discovery that the homozygous *Gpdh1* mutant stock used to study loss of the maternal GPDH1 enzyme pools exhibited striking differences in the steady state levels of amino acids and tricarboxylic acid (TCA) metabolites when compared with the control strain and F1 mutant generation. These unexpected findings provide insight towards understanding how *Drosophila* metabolism is rewired to support the complete loss of a major enzyme involved in central carbon metabolism.

## METHODS

### Drosophila melanogaster husbandry and genetics

Fly stocks were maintained on Bloomington *Drosophila* Stock Center (BDSC) food at 25°C. The *Gpdh1*^*A10*^ mutant strain was described in a previous study from our lab (Li *et al*. 2019). For all the experiments, 50 adult virgins were mated with 25 males and the embryos were collected on molasses agar plates with yeast paste for 4 hours, as previously described (Li and Tennessen 2017). *Gpdh1*^*A10*^ zygotic mutant larvae were collected by crossing *Gpdh1*^*A10*^*/CyO, P{w[+mC]=GAL4-twi*.*G}2*.*2, P{w[+mC]=UAS-2xEGFP}AH2*.*2* males and females and selecting for larvae lacking GFP expression. The *Gpdh1* maternal-zygotic mutant stock was established by crossing *Gdph1*^*A1*0^ zygotic mutant males and females.

### Ovaries dissection and imaging

Ovaries were dissected from *Gpdh1*^*A10/+*^, *Gpdh1*^*A10*^ *(Z) and Gpdh1*^*A10*^ *(M/Z)* females that were raised on yeast. Ovaries were fixed with 4% formaldehyde, rinsed twice with phosphate buffer saline (PBS, pH 7.4), stained with DAPI for 30 minutes, and mounted on slides using vector shield with DAPI (Vector Laboratories; H-1200-10). Slides were imaged using a Leica SP8 confocal microscope.

### Fecundity analysis

Twenty virgin females from *Gpdh1*^*A10/+*^ (control), *Gpdh1*^*A10*^ (Z) and *Gpdh1*^*A10*^ (M/Z) were mated with ten *w*^*1118*^ males in a small mating chamber. The total number of eggs laid during a two-hour period was quantified for each mating chamber.

### Viability assays

Embryonic viability of *Gpdh1*^*A10/+*^, *Gpdh1*^*A10*^ *(Z)*, and *Gpdh1*^*A10*^ *(M/Z)* genotypes were conducted by crossing 50 female virgins with 25 males in an egg-laying bottle that contained a molasses agar plate partially covered with yeast paste (see Li and Tennessen 2017). The egg-laying plate was replaced every 24 hours for two days, after which time a fresh egg-laying plate was placed in the bottle and eggs were collected for two hours. The egg-laying plate was removed, and following an 8 hour incubation at 25°C to allow for onset of GFP expression from the balancer chromosomes, thirty non-GFP embryos were identified by circling the surrounding agar with a dissecting needle. The number of eggs that hatched to become L1 larvae was recorded 24 hours later.

Larval viability was measured by placing 20 synchronized embryos of each genotype on molasses agar plates with yeast paste and measuring time until pupariation. Pupae were subsequently transferred into a glass vial containing BDSC food and monitored until eclosion.

### Longevity assay

Virgin female and male adults of the indicated genotypes were separated into glass vials (n = 10 adults/vial) and maintained at 25°C. Flies were transferred to fresh vials every 5 days without the use of carbon dioxide. The number of dead adults in each vial was recorded daily.

### Fat body staining

Mid-L2 larval fat bodies were fixed and stained as previously described (Tennessen *et al*. 2014a). Briefly, fat bodies were fixed with 4% formaldehyde, rinsed twice with PBS, once with 50% ethanol, and then stained for 2 minutes at room temperature using filtered 0.5% Solvent Black 3 (CAS Number 4197-25-5; Sigma 199664) dissolved in 75% ethanol. Samples were sequentially rinsed with 50% ethanol, 25% ethanol, and PBS. Stained tissues were mounted on a microscope slide with vector shield with DAPI (Vector Laboratories; H-1200-10).

### Gas Chromatography-Mass Spectrometry (GC-MS) analysis

GC-MS analysis was performed at the Indiana University Mass Spectrometry Facility as previously described (Li and Tennessen 2018). All samples contained 25 mid-L2 larvae and six biological replicates were analyzed per genotype. GC-MS data was normalized based on sample mass and an internal succinic-d4 acid standard that was present within the extraction buffer. Data were analyzed using Metaboanalyst version 5.0 following log transformation (base 10) and Pareto Scaling (Pang *et al*. 2021).

### Adult body mass measurements

Ten adult male or female flies were collected one day post-eclosion, placed into pre-tared 1.5 mL microfuge tubes, and the mass was measured using a Mettler XS 105 weighing analytical balance. Six independent samples were measured per genotype.

### Statistical Analysis

Unless noted, statistical analysis was conducted using GraphPad Prism v9.1. Data are presented as scatter plots, with error bars representing the standard deviation and the line in the middle representing the mean value. Unless noted, data was compared using a Kruskal-Wallis test followed by a Dunn’s multiple comparison test. Longevity data was analyzed using a log-rank (Mantel-Cox) test.

## RESULTS AND DISCUSSION

### GPDH1 is required for oogenesis and embryonic viability

Previous studies indicate that GPDH1 is expressed in the ovary and maternally loaded into the egg (Wright and Shaw 1969; Wright and Shaw 1970; Casas-Vila *et al*. 2017). Considering that homozygous *Gpdh1* mutant strains are reported to be sub-viable (O’brien and macIntyre 1972b; O’brien and Shimada 1974; Kotarski *et al*. 1983; Merritt *et al*. 2006), we examined the possibility that GPDH1 serves essential roles during early development that have been previously overlooked. As a first step towards testing this possibility, we dissected the ovaries from the homozygous *Gpdh1*^*A10*^ mutant flies – both from F1 mutants (generated by crossing *Gpdh1*^*A10*^*/CyO, twi-GFP* parents and selecting for GFP^-^ larvae), as well as from a homozygous *Gpdh1*^*A10*^ mutant strain (designated *M/Z* to indicate a complete absence of enzyme during development). We found that the ovaries from both F1 *Gpdh1*^*A10*^ females and *Gpdh1*^*A10*^ *M/Z* females were smaller than those observed in control females and contained fewer late-stage ovarioles (Figure 2A-C). Notably, we regularly observed degenerating egg chambers in mutant embryos (See Figure 2B, arrow), indicating that loss of GPDH1 in females limits oocyte production. Consistent with the observed egg chamber defects, both classes of *Gpdh1* mutants exhibit significant decreases in egg-laying as compared with controls (Figure 2D).

**Figure 2.**
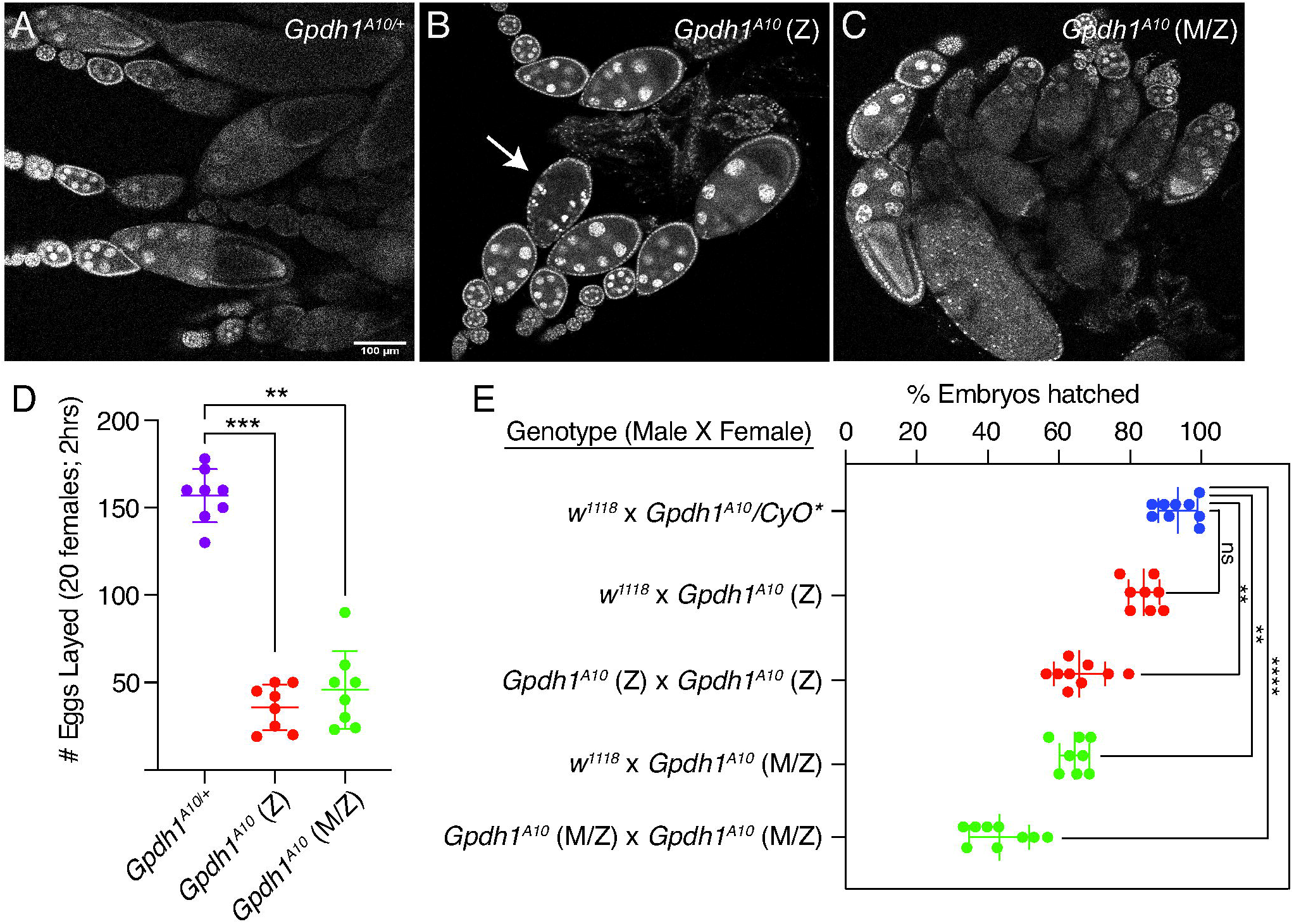
*Gpdh1* is required for oocyte development and embryonic viability. (A-C) Ovaries were dissected from 3-day old females and stained with DAPI. When compare to the *Gpdh1*^*A10/+*^ control strain (A), the ovaries of (B) F1 generation *Gpdh1*^*A10*^ mutants and (C) *Gpdh1*^*A10*^ *M/Z* mutants display (B,C) fewer later stage oocytes and (B) degenerating egg chambers (see arrow). (D) The fecundity of F1 generation *Gpdh1*^*A10*^ mutants and *Gpdh1*^*A10*^ *M/Z* mutants are significantly decreased when compared with controls. (E) Embryos produced by either F1 generation *Gpdh1*^*A10*^ mutants or *Gpdh1*^*A10*^ *M/Z* mutants exhibit significant mortality when compared with the controls and independent of paternal genotype. \**P*< 0.05. \*\**P*< 0.01. \*\*\**P*< 0.001. *P*-values calculated using a Kruskal-Wallis tests followed by a Dunn’s test.

As a complement to our studies of oogenesis, we also examined if the maternal and zygotic GPDH1 enzyme pools are required for embryonic development. Females of a control strain (*Gpdh1*^*A10/+*^), F1 *Gpdh1* mutants (*Gpdh1*^*A10*^) and the *Gpdh1* mutant strain (*Gpdh1*^*A10*^ *M/Z)* were crossed with either *w*^*1118*^ controls or *Gpdh1*^*A10*^ mutant males and the resulting offspring were scored for the percentage of embryos that hatched to first instar larvae (L1). We observed that embryos from *Gpdh1* mutant mothers of either genetic background died at a significantly higher rate than those produced by heterozygous control mothers, regardless of the parental genotype (Figure 2E), indicating that both the maternal and zygotic GPDH1 pools are required during embryogenesis. Overall, our findings that *Gpdh1* mutants display oogenesis defects and embryonic lethality explains why *Gpdh1* mutant females exhibit such low levels of fecundity when compared with control strains.

### Zygotic and Maternal-Zygotic *Gpdh1* mutants display similar defects during larval development and adult longevity

The observed requirement for maternal GPDH1 in embryogenesis led us to reevaluate if loss of this enzyme pool also influences development and lifespan in later life stages. Like previous reports of *Gpdh1* zygotic mutants, we found the *Gpdh1 M/Z* mutant larvae develop more slowly than heterozygous controls and ∼20% fail to initiate metamorphosis (Figure 3A). These developmental delay phenotypes, however, are indistinguishable from *Gpdh1* zygotic mutants (Figure 3A). Moreover, *Gpdh1 M/Z* mutants eclose at the same rate as zygotic mutants (Figure 3B). The only observable difference between the zygotic and *M/Z* mutants is that newly eclosed *Gpdh1 M/Z* mutants females exhibit slightly increased body mass (Figure 3C), however, we are unable to rule out the possibility that this phenotype is due to differences in genetic background.

**Figure 3.**
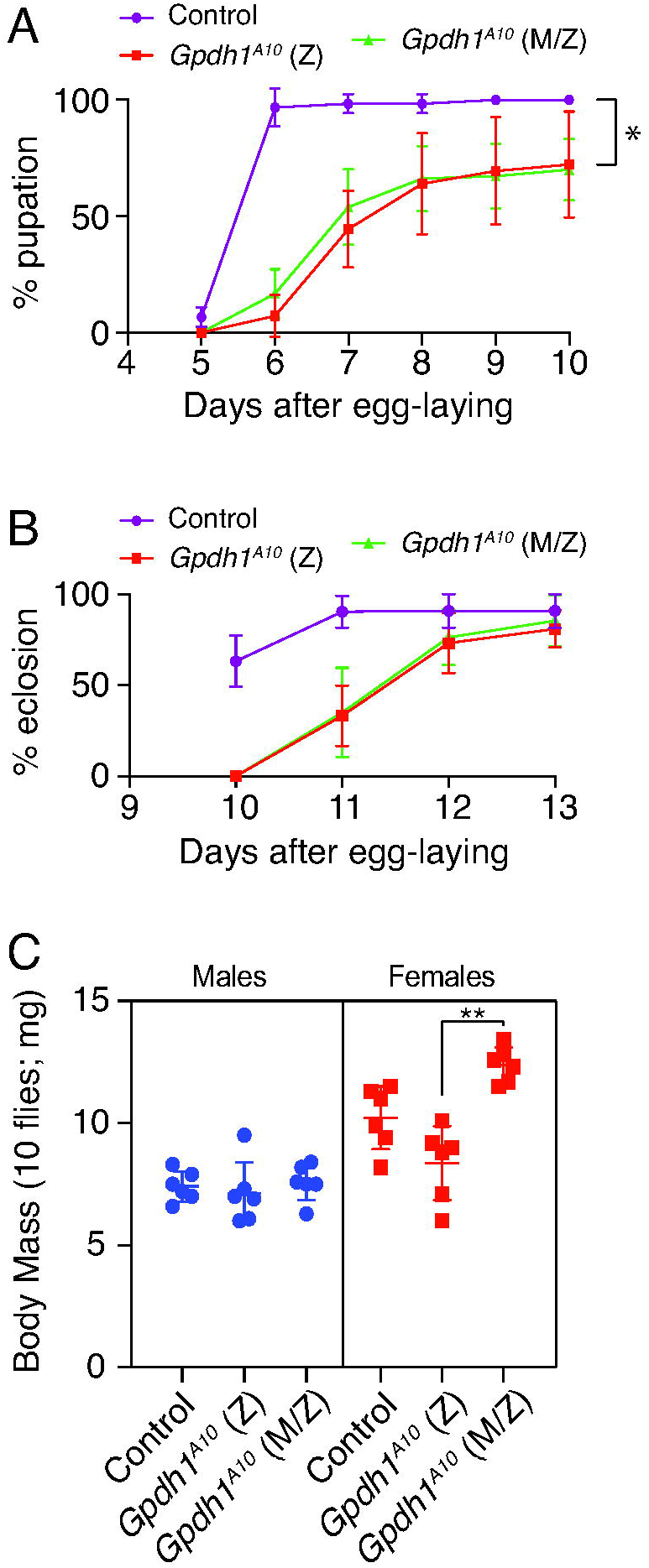
*Gpdh1* zygotic and *Gpdh1* maternal-zygotic mutants exhibit similar developmental delays. *Gpdh1*^*A10/+*^ controls, F1 generation *Gpdh1*^*A10*^ mutants, and *Gpdh1*^*A10*^ *M/Z* mutants were analyzed for (A) time from egg-laying to pupariation, (B) time from egg-laying to eclosion, and (C) body mass one day after eclosion. For (B), the % eclosion value represents the percent of pupae that successfully eclosed and does not include the 20% of larvae that failed to pupariate. \*\**P*< 0.01. *P*-values calculated using a Kruskal-Wallis tests followed by a Dunn’s test.

As a complement to our studies of larval development, we also examine the viability of *Gpdh1* mutant adults. Here too, we observe little difference between F1 *Gpdh1* mutants and *M/Z* mutants. Consistent with earlier findings (Merritt *et al*. 2006), we observed that both the *Gpdh1*^*A10*^ zygotic mutant females and *M/Z* mutant females are short-lived when compared with a heterozygous controls strain, with a majority mutant animals dying within two weeks of eclosion (Figure 4A). However, we observed that the short lifespan phenotype was milder in F1 *Gpdh1*^*A10*^ mutant males and absent in the *Gpdh1*^*A10*^ *M/Z* mutant males (Figure 4B). The observed decrease in female *Gpdh1* lifespan is apparent while maintaining the maternal-zygotic stock on Bloomington *Drosophila* Stock Center food. Although bottles of the *Gpdh1*^*A10*^ *M/Z* mutant strain contains approximately equal ratios of males:females two days after eclosion, this ratio becomes significantly skewed two weeks post-eclosion due to females dying earlier than male mutants (Figure 4C). Overall, our findings reveal that nearly all growth, development, and viability phenotypes exhibited by the *Gpdh1*^*A10*^ *M/Z* mutants are no more severe than those observed in the F1 *Gpdh1*^*A10*^ mutants.

**Figure 4.**
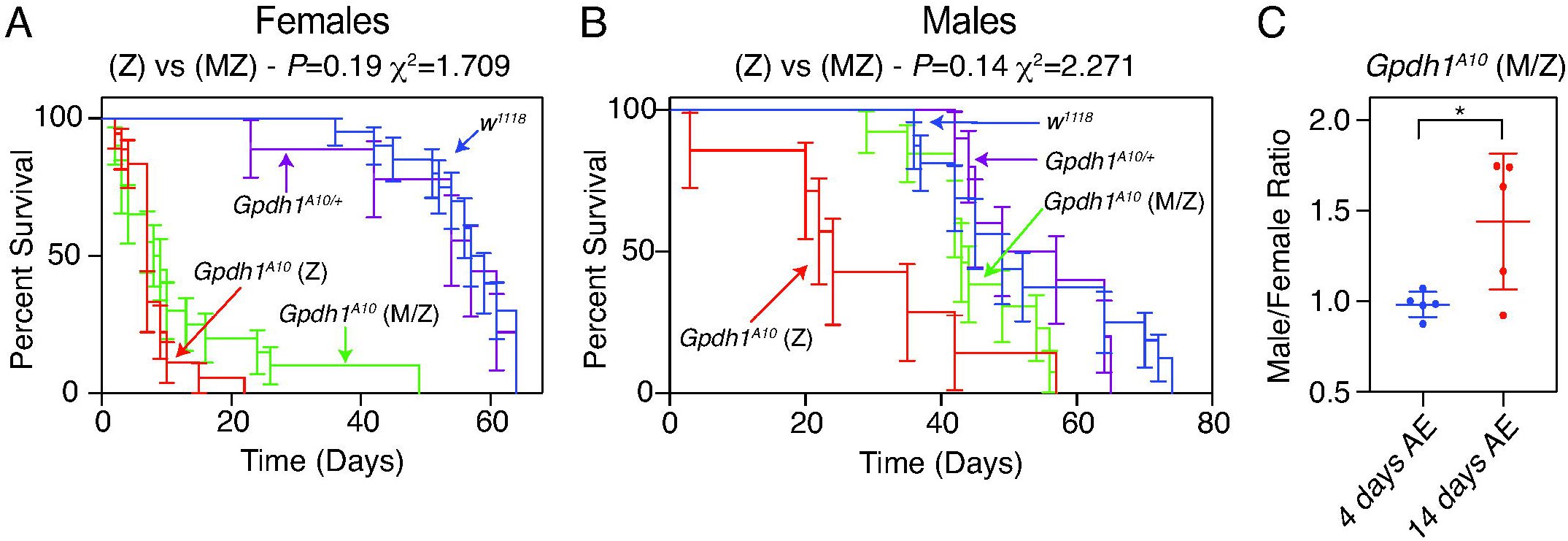
*Gpdh1* mutant females are short-lived. The lifespan of (A) females and (B) males from *Gpdh1*^*A10/+*^ heterozygous controls, F1 generation *Gpdh1*^*A10*^ mutants, and a *Gpdh1*^*A10*^ *M/Z* mutant strain were analyzed for 80 days after eclosion. Longevity data was analyzed using a log-rank (Mantel-Cox) test. (C) Ratio of males to females in culture bottles either 2 or 14 days after eclosion. \**P*< 0.05. *P*-values calculated using an unpaired t-test.

### *Gpdh1* zygotic mutants and maternal-zygotic mutants exhibit significant differences in amino acid metabolism

Larval development is able to compensate for loss of zygotic GPDH1 by inducing compensatory changes in central carbon metabolism, rendering *Gpdh1* mutants dependent on LDH and GPO1 activity while also inducing significant changes in redox balance and steady-state amino acid levels (Davis and macIntyre 1988; Li *et al*. 2019). These earlier metabolic studies were conducted by crossing males and females from a heterozygous *Gpdh1*^*-*^*/CyO,twi-GFP* stock and selecting for homozygous offspring, thus providing a readout of how loss of zygotic GPDH1 affects larval metabolism. In this regard, our homozygous *Gpdh1 M/Z* mutant strain provides a unique opportunity to determine if loss of both the maternal and zygotic enzyme pools induce a distinct metabolic profile when compared with zygotic mutants. Towards this goal, we first examined larval triglyceride (TAG) levels in the heterozygous control, F1 *Gpdh1* mutant, and the homozygous *Gpdh1 M/Z* strain. Consistent with previous studies (Merritt *et al*. 2006), we observed similar decreases in fat body TAG levels of the *Gpdh1*^*A10*^ zygotic and *Gpdh1*^*A10*^ *M/Z* mutant larvae when compared with *Gpdh1*^*A10/+*^ heterozygous controls (Figure 5A-C). These results indicate that loss of the maternal GPDH1 pool does not exacerbate the *Gpdh1* TAG mutant phenotype.

**Figure 5.**
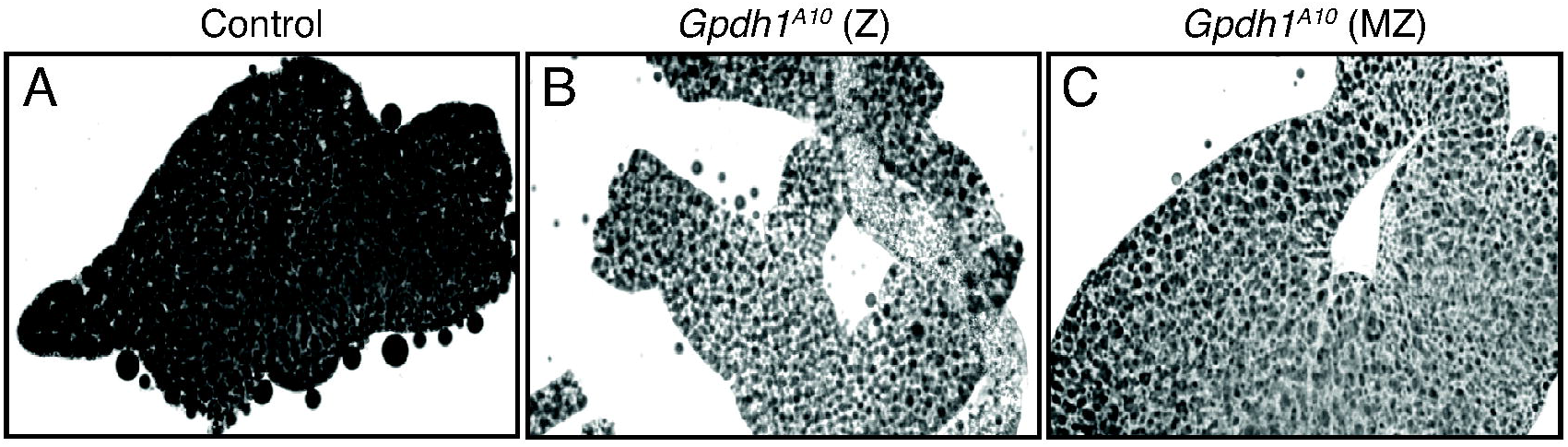
TAG levels are significantly decreased in *Gphd1* mutants. Representative images of mid-L2 fat bodies stained with Black Solvent 2 and imaged using bright field microscopy. When compared with (A) heterozygous *Gpdh1*^*A10/+*^ control, both (B) F1 generation *Gpdh1*^*A10*^ mutants (C) a *Gpdh1*^*A10*^ *M/Z* mutant strain exhibit a similar decrease in lipid levels.

We next used a targeted GC-MS-based approach to compare the levels of amino acids, TCA cycle intermediates, and glycolytic end products in mid-L2 larvae of *Gpdh1*^*A10/+*^ heterozygous controls, *Gpdh1*^*A10*^ zygotic mutants, and *Gpdh1*^*A10*^ *M/Z* mutants (Table S1). When the resulting metabolomics data was analyzed using principal component analysis, the *Gpdh1*^*A10*^ *M/Z* mutant samples clearly separated from both the *Gpdh1*^*A10*^ zygotic mutants and the *Gpdh1*^*A10/+*^ heterozygous controls (Figure 6A), suggesting that the three genotypes have distinct metabolic profiles. A closer analysis of the datasets revealed that the changes observed in *Gpdh1*^*A10*^ zygotic mutant larvae mimic those observed previously (Li *et al*. 2019) – G3P levels were significantly decreased in relative to the control strain (Figure 6B,C) and both lactate and 2-hydroxyglutarate remained at comparable levels (Figure 6B, D, E). The relative abundance of some amino acids and TCA cycle intermediates were also decreased in the zygotic *Gpdh1* mutant larvae compared with *Gpdh1*^*A10/+*^ controls (Figure 6B,F,G,H).

**Figure 6.**
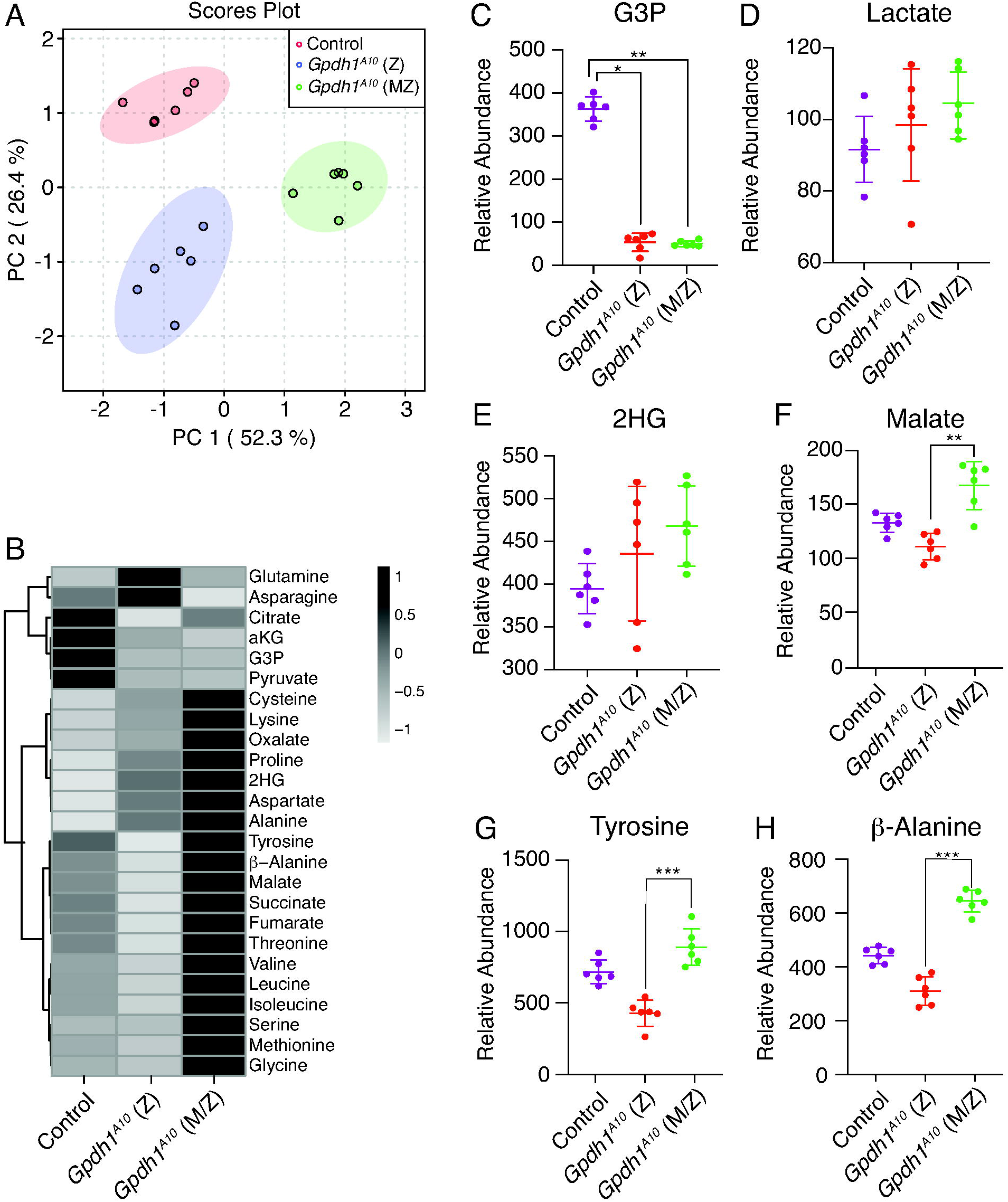
Metabolomic analysis of *Gpdh1* zygotic and *Gpdh1* maternal-zygotic mutants. A targeted GC-MS-based metabolomics method was used to compare the relative abundance of G3P, lactate, 2HG and amino acids, between heterozygous *Gpdh1*^*A10/+*^ controls, *Gpdh1*^*A10*^ zygotic mutants and *Gpdh1*^*A10*^ maternal-zygotic mutants. (A) PCA plot showing that the control, F1 generation *Gpdh1* mutants *(Z)*, and the *Gpdh1 M/Z* mutant strain separate clearly in their metabolomic profile. (B) Heatmap showing the increase in the relative abundance of amino acids in *Gpdh1 M/Z* mutants when compared to the F1 generation *Gpdh1* zygotic mutants and the *Gpdh1*^*A10/+*^ controls. (C-H) The relative abundance of (C) G3P, (D) lactate, and (E) 2HG, (F) malate, (G) tyrosine, and (H) β-alanine are represented as scatter plots with the horizontal lines representing the mean value and standard deviation. \**P*< 0.05. \*\**P*< 0.01. \*\*\**P*< 0.001. *P*-values calculated using a Kruskal-Wallis tests followed by a Dunn’s test. Analysis in (A) and (B) conducted using MetaboAnalyst 5.0 (see methods).

While the metabolite changes observed in *Gpdh1* zygotic mutants largely confirmed previous studies, the metabolic profile of the *Gpdh1*^*A10*^ *M/Z* mutants exhibited striking differences. Even though G3P, lactate, and 2-hydroxyglutare levels were comparable between the two mutant genotypes (Figure 6B-E), many of the same amino acids that were either decreased or unchanged in the zygotic mutants were significantly elevated in the *M/Z* mutant background (Figure 6B, F-H). For example, relative to the control strain, tyrosine and β-alanine were decreased by 40% and 30%, respectively, in zygotic mutants but increased by 40% and 25% in the maternal-zygotic mutant (Figure 6G,H). A similar trend was observed with the TCA intermediates succinate, fumarate, and malate (Figure 6B), with the relative abundance of malate exhibiting a ∼15% decrease in zygotic mutants and a 25% increase in maternal-zygotic mutants when compared to the heterozygous control strain. The results are notable because they support a previously published hypothesis that *Gpdh1* mutants experience compensatory changes in malate metabolism (Merritt *et al*. 2006). Overall, our results reveal that a strain lacking both the maternal and zygotic GPDH1 enzyme pool exhibit changes in the steady-state levels of amino acid and TCA cycle intermediates that are opposite of those observed in mutants lacking only the zygotic enzyme contribution. Future studies should determine if these changes are due to either genetic selection when establishing the homozygous *Gpdh1* mutant stock or directly result from loss of the maternal GPDH1 enzyme pool.

## DISCUSSION

Here we demonstrate the *Drosophila* enzyme GPDH1 serves essential roles in both oogenesis and embryogenesis. Our findings expand the known roles for GPDH1 and raise questions as to what function this enzyme serves during early development. Considering that the purpose of GPDH1 activity differs in a context specific manner (*e*.*g*., TAG synthesis in fat body, ATP production in flight muscle), our findings raise the question as to the function of GPDH1 in these developmental contexts. Moreover, G3P levels are known to increase over the course of embryogenesis (Tennessen *et al*. 2014b), suggesting that this compound serves a unique role in the developing embryo.

Our findings also demonstrate that larvae lacking both maternal and zygotic GPDH1 activity exhibit developmental phenotypes that are largely indistinguishable from *Gpdh1* zygotic mutants. However, targeted metabolomics analysis indicates that *Gpdh1 M/Z* mutants exhibit metabolic phenotypes that are more severe than the zygotic mutants, raising the question as to how loss of maternal enzyme pool can influence the larval metabolic program in such a dramatic manner. While we can’t rule out the possibility that maternal GPDH1 activity establishes a metabolic state in the embryo that persists into larval development, a more likely explanation stems from the fact that oogenesis and embryogenesis are energetic processes that impose intense demands on cellular metabolism. As a result, generation of the homozygous *Gpdh1* mutant strain would inevitably select for background mutations that compensate for loss of GPDH1 activity in the ovary and embryos – a hypothesis that is supported by previous observations. For example, even though GPDH1 is essential for maintaining ATP production in flight muscle (O’brien and macIntyre 1972b; Wojtas *et al*. 1997; Merritt *et al*. 2006), *Gpdh1* mutants slowly regain the ability to fly when maintained in lab culture (O’brien and Shimada 1974), indicating that other metabolic processes must be able to compensate for loss of GPDH1 activity. Similarly, *Gpdh1* larvae only exhibit slight developmental delays despite displaying a significant disruption in redox balance. This ability of larvae to maintain a somewhat normal growth rate in the absence of GPDH1 depends on the enzymes LDH and GPO1, as loss of either enzyme in a *Gpdh1* mutant background enhances the mutant phenotype (Davis and macIntyre 1988; Li *et al*. 2019). These earlier studies, combined with our findings and previous observations that *Gpdh1* phenotypes are highly dependent on genetic background (Merritt *et al*. 2006), indicate that GPDH1 functions within a complex and highly adaptable metabolic network that warrants further study.

Regardless of the reason for why *Gpdh1* maternal-zygotic mutants exhibit significant metabolic differences when compared with zygotic mutants, our study highlights a poorly understood relationship between G3P and amino acid metabolism. While we are unsure as to the significance of this metabolic relationship in the fly, our findings are consistent with studies of a mouse model of GPD1 deficiency, which induces compensatory amino acid metabolism during fasting in mice (Sato *et al*. 2016). One intriguing possibility is that the *Gpdh1* maternal-zygotic mutants induce changes in one of the central metabolic regulating pathways that control amino acid metabolism. In this regard, the amino acid sensor Tor is not only known to regulate the yeast glycerol-3-phosphate dehydrogenase 1 activity in yeast (Lee *et al*. 2012), but the GPDH1 substrate, DHAP (see Figure 1), but also activates Tor in mammalian cell culture (Orozco *et al*. 2020). These correlations between Gpdh1 and Tor should be the subject of future investigations.

Our studies also revealed changes in the relative abundance of a subset of TCA acid cycle intermediates. These observations suggest that mitochondrial metabolism in *Gpdh1* maternal-zygotic mutants is fundamentally altered when compared with controls and zygotic mutants. Considering that GPDH1 is an essential enzyme in the G3P shuttle, which shuttles electrons to the mitochondria for ATP production (Figure 1), the observed increases in succinate, fumarate, and malate hint at the possibility that maternal-zygotic mutants exhibit significant changes in mitochondrial metabolism. However, since we observe no changes in the relative abundance of alanine, lactate, and 2-hydroxyglutarate, which are commonly elevated in flies that experience disruption of mitochondrial activity (Feala *et al*. 2007; Coquin *et al*. 2008; Campbell *et al*. 2019; Mahmoudzadeh *et al*. 2020), the significance of these changes in *Gpdh1* maternal-zygotic mutant remain unclear.

Overall, our results again emphasize that GPDH1 serves a unique role in *Drosophila* metabolism. The studies presented herein both confirms a large body of literature regarding the role of GPDH1 in physiology, development, and lifespan and reveals new roles for this enzyme in oogenesis, embryogenesis, and amino acid metabolism. Moreover, considering that several studies hint at a key role for human GPD1 in cancer metabolism (Zhou *et al*. 2017; Rusu *et al*. 2019; Liu *et al*. 2021; Xia *et al*. 2021), our findings highlight the need to better understand how this highly studied enzyme influences gene expression, cell growth and differentiation, and metabolic signaling networks.

## ACKNOWLEDGMENTS

We thank the Bloomington *Drosophila* Stock Center (NIH P40OD018537) for providing fly stocks, the *Drosophila* Genomics Resource Center (NIH 2P40OD010949) for genomic reagents, and Flybase (NIH 5U41HG000739) and the Indiana University Light Microscopy Imaging Center. Targeted GC-MS analysis was conducted using instruments housed in the Indiana University Mass Spectrometry Facility, which is supported, in part, by NSF MRI Award 1726633. J.M.T. is supported by the National Institute of General Medical Sciences of the National Institutes of Health under a R35 Maximizing Investigators’ Research Award (MIRA; 1R35GM119557).

## SUPPLEMENTAL TABLES

**Table S1. GC-MS analysis of control, *Gpdh1* zygotic mutants, and *Gpdh1* maternal-zygotic mutants**. Samples contained 25 mid-L2 larvae. Data normalized to sample mass and a d4-succinic acid internal standard.

